# Substrate binding and specificity appear as major forces in the functional diversification of eqolisins

**DOI:** 10.1101/167544

**Authors:** María Victoria Revuelta, Nicolas Stocchi, Priscila Ailín Lanza Castronuovo, Mariano Vera, Arjen ten Have

**Affiliations:** Instituto de Investigaciones Biológicas (IIB-CONICET-UNMdP), Facultad de Ciencias Exactas y Naturales, Universidad Nacional de Mar del Plata, CC 1245, 7600 Mar del Plata, Argentina; QUIAMM-INBIOTEC-CONICET, Department of Chemistry – Facultad de Ciencias Exactas y Naturales, Universidad Nacional de Mar del Plata, Funes 3350, 7600 Mar del Plata, Argentina

## Abstract

**Background:** Eqolisins are rare acid proteases found in archaea, bacteria and fungi. Certain fungi secrete acids as part of their lifestyle and interestingly these also have many eqolisin paralogs, up to nine paralogs have been recorded. This suggests functional redundancy and diversification, which was the subject of the research we performed and describe here.

**Results:** We identified eqolisin homologs by means of iterative HMMER analysis of the NR database. The identified sequences were scrutinized for which we defined novel hallmarks, identified by molecular dynamics simulations of mutants of highly conserved positions, using the structure of an eqolisin that was crystallized in the presence of a transition state inhibitor. Four conserved glycines were shown to be required for functionality. A substitution of W67F is shown to be accompanied by the L105W substitution. Molecular dynamics shows that the W67 binds to the substrate via a π-π stacking and a salt bridge, the latter being stronger in a virtual W67F/L105W double mutant of the resolved structure of Scytalido-carboxyl peptidase-B (PDB ID: 2IFW)). Additional likely fatal mutants are discussed.

Upon sequence scrutiny we obtained a set of 233 sequences that in all likelihood lack false positives. This was used to reconstruct a Bayesian phylogenetic tree. We identified 14 putative specificity determining positions (SDPs) of which four are explained by mere structural explanations and nine seem to correspond to functional diversification related wit substrate binding ans specificity. A first sub-network of SDPs is related to substrate specificity whereas the second sub-network seems to affect the dynamics of three loops that are involved in substrate binding.

**Highlights:** Eqolisins are acid proteases found in prokaryotes and fungi only.

The recently co-evolved W67F-L105W substitutions promote substrate binding

Two Specificity Determining Networks, SDN1 and 2, were identified

SDN1 has four Specificity Determining Positions involved in substrate specificity

SDN2 has five Specificity Determining Positions involved in loop-substrate dynamics

## 1 Introduction

### 1.1 Acid Proteases

The three major families of acid or carboxyl peptidases recognized by MEROPS are the well studied, pepstatin-sensitive, eukaryotic aspartic proteinases (A01, APs, for review see [1]), part of the aspartic proteinase clan A; the more recently identified sedolisins (S53) and the also novel eqolisins or glutamic peptidases (G01), both recently reviewed [2]. Both sedolisins and eqolisins were first thought to be pepstatin-insensitive variants of APs but the structures that were resolved showed they are unrelated. The rather recent discovery of these enzymes means we have relatively little knowledge. However, since both sedolisins and eqolisins are typically active below pH 4, they are interesting both from a fundamental and an applied angle. Here we study the eqolisins by means of molecular evolution and dynamics, a study on sedolisins we recently reported elsewhere [3].

### 1.2 Eqolisins are Glutamic Peptidases

Eqolisins have been described in archaea, bacteria [4] and fungi but, interestingly, are not found in any non-fungal eukaryote [5]. Furthermore, eqolisins seem to have a rather unique and simple fold, which makes them straightforward targets for structure-function prediction. Eqolisins are endopeptidases synthesized as a preproprecursor proteins. The preprosegments are aproximately 55 amino acids in length, and the prosegments are rich in positively charged residues. Scytalido-carboxyl peptidase-B (SCP-B) from *Scytalidium lignicolum* is the enzyme that first described the eqolisin family of peptidases and its structure has been resolved (PDB ID: 2IFW) [6]. Point mutation analyses revealed its catalytic site is formed by a catalytic dyad (Q53, E136), hence the name EQolisin. SCP-B has a narrow substrate specificity, with preference for small, basic residues, rather than the typical hydrophobic residues preferred by APs [1]. D43 is a residue that is important for SCP-B structure since the D43A mutant has approximately 20% of the original activity [7] [8]. Other characterized eqolisins are *Aspergillus niger* carboxyl peptidase (ANCP) [9], which 3D structure has also been resolved (1Y43) [10], *Sclerotinia sclerotiorum* carboxyl peptidase [11], *Cryphonectria parasitica* peptidases B and C [12], *Talaromyces emersonii* carboxyl peptidase TGP1 [13] and bacterial *Alicyclobacillus* sp pepG1 [4].

Eqolisins are composed of two seven-stranded anti-parallel β-sheets that fold in parallel and bend to form a structure resembling a half-pipe that forms the binding cleft (See Fig. 1). The catalytic Q53 and E136 (numbering according to SCP-B structure 2IFW) stick into the binding cleft. Pillai [14] described the 70’s loop (Tyr71-Gly80) and the β-loop (Cys141-Cys148) that, upon interaction with a transition state inhibitor appear to move inwards and, thus, likely play a role in substrate binding and catalysis. They also indicated a high structural similarity of eqolisins with the carbohydrate-binding concanavalin A-like lectins/glucanases superfamily. Additional analyses indicate that residues Y64 to Y71 are highly conserved across all members of the G1 family [14].

**Fig. 1.**
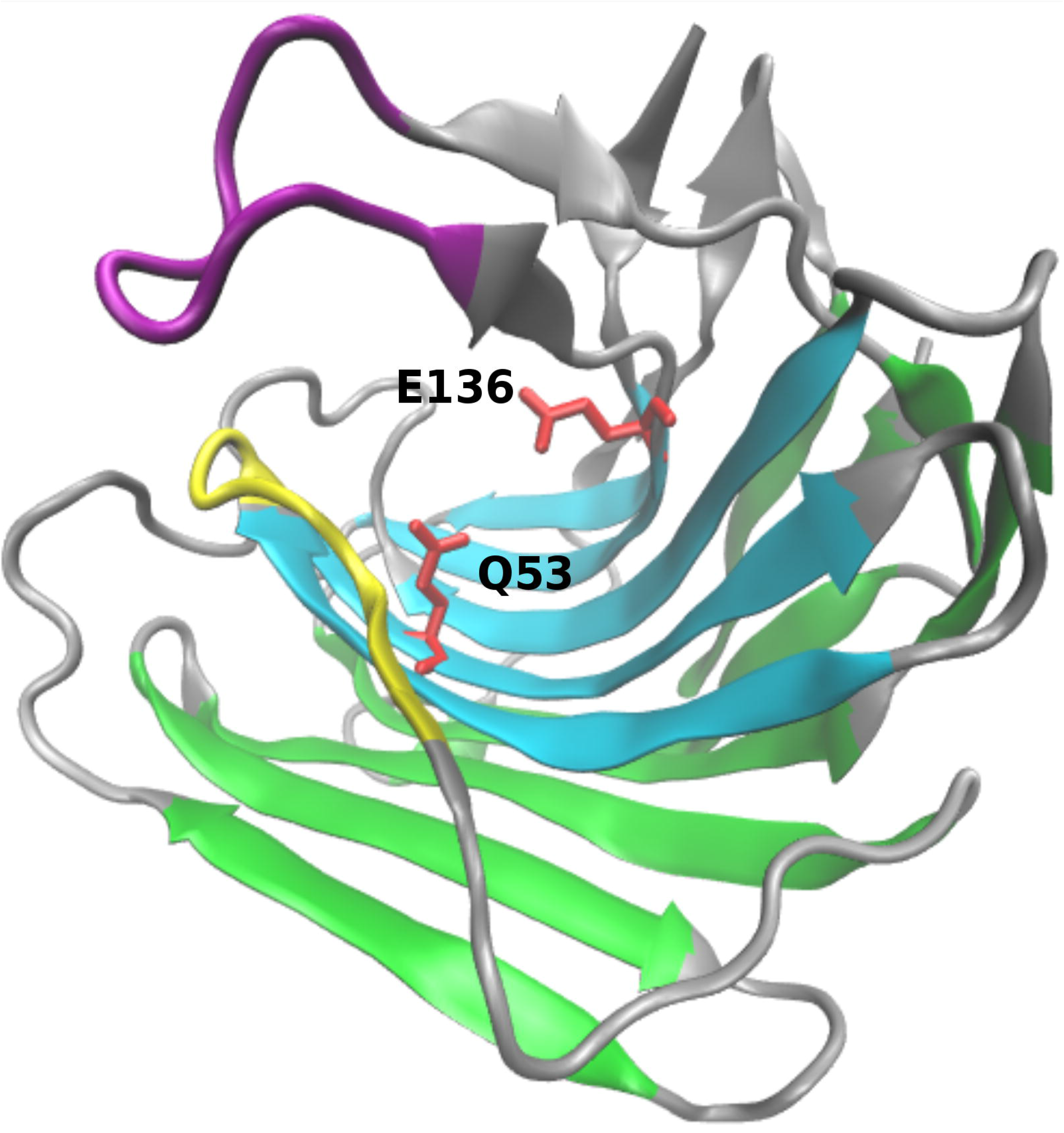
Structure of SCP-B Eqolisin. Eqolisins consist of two pleated β-sheets (Cyan and green for inner and outer sheet, respectively) that fold into a double convex halfpipe. E136 and Q53 (red licorice) stick into the binding cleft and form the catalytic site. The β-loop (purple) and 70’s-loop (yellow) have been shown to migrate inwards during substrate binding.

### 1.3 Fungal Eqolisins have undergone a Process of Functional Redundancy and Diversification.

Since eqolisins have only been recently described, not much is known in terms of biological function. *T. emersonii*’s eqolisin gene *tgp1* was shown to be induced in the presence of an extracellular protein source, displays a rather broad specificity, is the most abundant protease in its secretome and is essential for the fungal growth [13]. Poussereau and collaborators described the importance of an eqolisin from plant pathogen *S. sclerotiorum* in the sunflower cotyledon infection process, suggesting a role in pathogenesis [11]. Closely related plant pathogenic fungus *Botrytis cinerea* contains nine paralogues, which suggests functional redundancy and diversification. This is also suggested by the fact that *B. cinerea* and *S. sclerotiorum,* as well as for instance many *Aspergilli* that seem to have relatively many eqolisins, secrete acids as part of their lifestyle. Interestingly, these also show relatively few basic subtilisins. Recently we studied the sedolisins [3], here, we explore the eqolisin protein family in order to identify which residues are likely required for function and which could be involved in functional diversification.

## 2 Materials and Methods

### 2.1 Identification of Eqolisin Homologues

The seed HMMER profile was made by means of *hmmbuild* using default settings and the MEROPS [15] G1 alignment of holozymes (HMMER Version 3.0 [16]). This was used to iteratively screen the 107 complete proteome dataset previously used to identify Aspartic Proteases [17] and the HMMER NR database (W1). All sequences identified with an E-value smaller than the HMMER exclusion threshold were considered as Eqolisin homologues. Upon data acquisition, sequences were scrutinized for the presence of catalytic residues Q53 and E136, and secondary structure elements according to available resolved structures 2IFW and 1Y43. The novel dataset was then aligned by MAFFT [18] and used for iteration of *hmmbuild* and *hmmsearch* at the HMMER website. Iterations were performed until data convergence. Finally, sequences with long (>15 aa) inserts that appeared to lack homologous counterparts in any of the collected sequences or in a homologous sequence identified by BLAST in the non redundant database of NCBI, were removed

### 2.2 MSA and Phylogenetic Analysis

Multiple protein sequence alignments (MSAs) were performed using MAFFT [18] with slow iteration mode. Trimming for phylogeny was performed with BMGE [19] using the command options *-t AA -m BLOSUM30 -b 1 -h 0.9*. This setting did not remove subsequences corresponding to major secondary structure elements. A maximum likelihood (ML) phylogeny was built using PhyML-a-bayes □ [20] with the WAG model, estimated proportion of invariable sites, four rate categories, estimated gamma distribution parameter and optimized starting BIONJ tree, as determined by a prior estimation using ProtTest [21], with 10000 bootstraps branch support measure. The ML tree was used as starting tree for Bayesian analysis using MrBayes [22]. Chains were initiated with ten perturbed trees using default settings until convergence was reached. Convergence was tested when split frequency was below 0.01 using Awty [23]. The resulting phylogenetic trees were viewed and edited with iTol version 2.0 [24] or Dendroscope [25].

### 2.3 Molecular Modelling and Dynamics

Models for static analysis were made by I-Tasser [26].

The right protonation state of the wild type (WT) structural model (PDB ID: 2IFW) was determined by 19 reasonable protomers at pH=4.1, selecting the one with the clearly smaller RMSD after 40 ns of simulation (see below). The mutants W67F, W67F-L105W, L105W, P72K and G8A-G41A-G44A-G55A (hereafter GAx4) were prepared by replacing the side chain in the WT followed by local minimization using the AMBERTools [27] LEAP facility. The general setup for the MD simulations were as follows: I) 2500 steps steepest descent minimization of whole system, keeping the protein positionally restrained and embedded into a box of TIP3P water molecules with a minimum distance of 10 Å to each wall, and Cl- or Na+ counterions to neutralize. II) 2500-5000 conjugate gradient minimization of the whole systems. III) 150 ps slowly heating in the NTV ensemble. IV) 40 ns of simulation in the NTP ensemble, at 1 atm and 298.15 K. The procedure III-IV was repeated in three independent trajectories using the Langevin and twice the Andersen termostat/barostats [28] [29]. Then, 40 ns of NTP simulation of the WT under Andersen thermostat/barostat were used as reference for further for cross-correlations analysis, lowest normal modes visualization, and salt- and H-bridges inspection and comparison with the results obtained for the four mutants discussed below. In the equilibrated system, the density slightly fluctuated around 1.019 g/mL. Electrostatic interactions were computed using the Particle Mesh Ewald (PME) method with a cutoff of 10 Å [30] [31]. Bonds involving hydrogen atoms were constrained using the SHAKE algorithm [31], this allowing for an integration time step of 0.0015 ps. The integration was done usign the pmemd.CUDA module of the AMBER14 program, with the ff14SB force field [32] [33]. For the inhibitor, since most “residues” were of proteic nature, the same force field was applied, by manually modifying the connectivity of its backbone, the residues were labeled with three capital letter for distinguishing them from the protein, starting from 207 to 213: ACE207, PHE208, LYS209, PHE210, PSA211, LEU212 and AAR213. The trajectories were analyzed using standard AMBER analysis tools. The criteria for analyzing the persistence of H-bonds were set to a maximum length of 3.2 Å (between the heavy atoms) and a maximum angle of 120º (donor-H-acceptor). Analysis of the hydrogen bonds, contacts persistency, mobility factors and cross correlation functions were done using ccptraj (AMBERTools 15 utilities) and VMD1.9.7 [34] which was also used for graphics rendering. Essential normal modes were calculated using the last 30-35 ns and processing with principal components analysis as implemented in ProDy (W2).

The free energy calculations were done using the MM-PBSA module of AMBER 14 and reported for the two models applied, *i. e.* Poisson-Boltzman (PB) and Generalized Born (GB) [35]. [36]]. The energetic analysis were done from 5.0 to 40.0 ns of simulation.

### 2.4 Additional Biocomputational Analyses

The SignalP [37] server was used to predict the presence of signal peptides, using complete sequences. Sequence LOGOS were created with WebLogo [38], and structures were aligned VMD [34] using the STAMP [39] extension.

## 3 Results and Discussion

### 3.1 Sensitive Identification of Eqolisin Homologues likely lacks Specificity

The identification of eqolisin encoding sequences was performed with HMMER [16] in order to obtain high sensitivity. A profile made from the holotype sequences of MEROPS [15] was used as a seed. Initially, a collection previously used for a study of aspartic proteases [17], of mostly fungal, but also other taxonomically well distributed eukaryotic complete proteomes were used. This resulted in few sequences, hence, the NR database was scanned and all identified sequences were combined and searched iteratively until data convergence. Then, since eqolisins have been described only recently and appear restricted to fungi, bacteria and a few archaea, there is little knowledge on which residues other than the catalytic Q53 and E136 are required. The initial sequence mining was therefore not specific and might have resulted in the inclusion of sequences from non-functional homologs (NFHs) or incorrectly predicted gene models. Such sequences would introduce noise in a direct but also an indirect way by negatively affecting the quality of the MSA. Given the broad taxonomic distribution we reasoned that residues that appear highly conserved, defined as >95% using Genedoc’s default scheme of conserved groups, in a preliminary MSA, must be important and might even be required. Structural analysis of apparent rare substitutions might shed light on eqolisin functionality and clarify which of the highly conserved residues are actually strictly conserved, thereby identifying NFHs.

Fig. 2. shows an excerpt of a preliminary MSA that, besides 13 strictly conserved positions, had 10 highly conserved (>95%) positions that were all studied as part of a rigid sequence scrutiny. The complete preliminary MSA with all mentioned sequences can be found in S datafile 1. In order to determine if the highly conserved positions, are indicative for additional strictly required amino acids among functional eqolisins, we performed molecular dynamics.

**Fig. 2.**
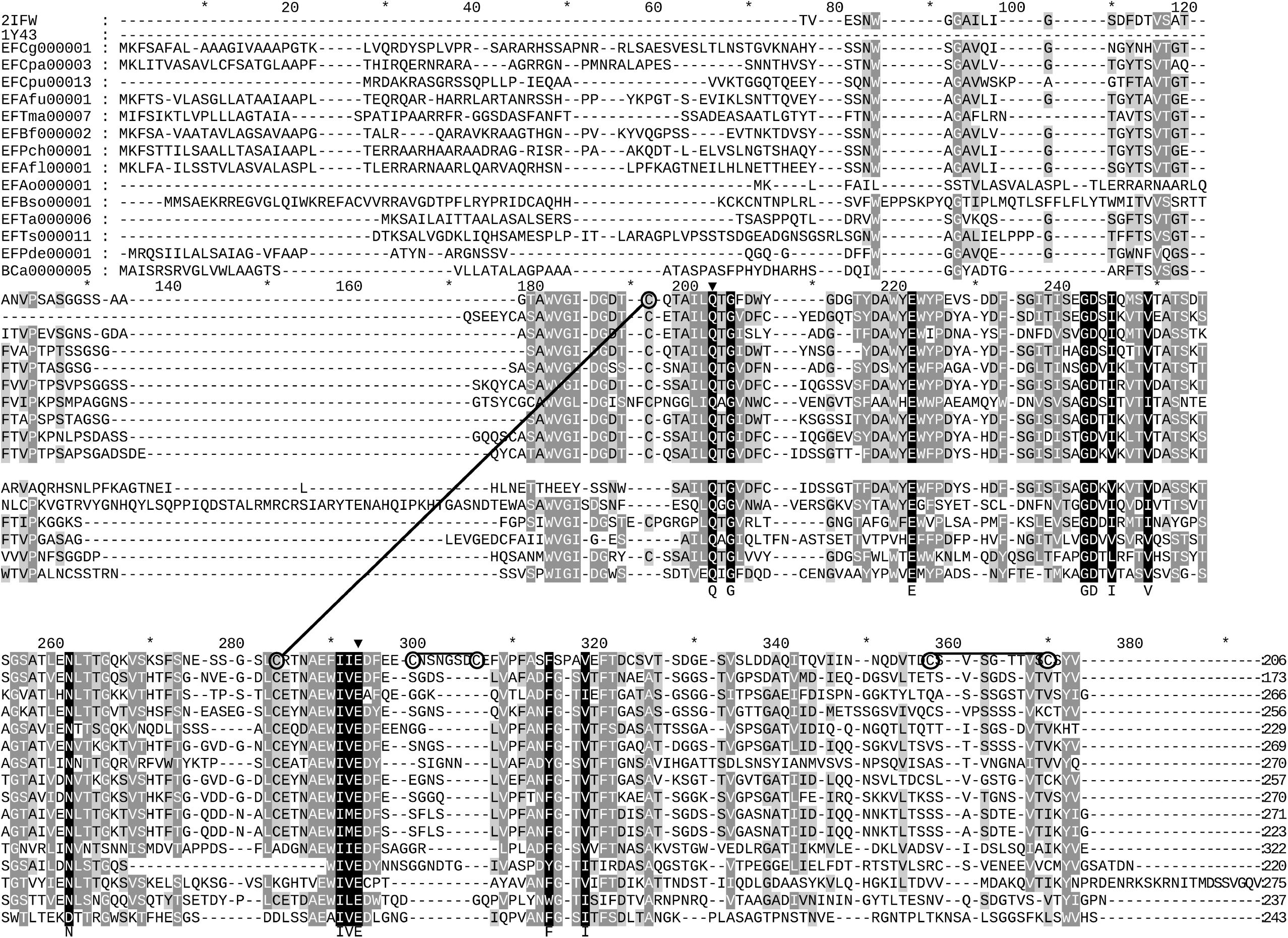
Excerpt of Multiple Sequence Alignment Eqolisins. Indicated are the sequences for which a structure has been resolved (PDB IDs 2IFW, 1Y43) as well a number of sequences selected to show overall sequence variation. The lower block contains sequences that were removed upon sequence scrutiny as described in the main text. The rulers indicate positions of hallmark residues and or positions described under the sequence scrutiny. Encircled cysteines are involved in disulfide bridges, indicated by lines. Shading indicates conservation (100% black; 80% dark-gray, 60% light-gray).

### 3.2 Wild Type Dynamics

We first determined the most optimal protonation state. The most likely protomer at pH=4.1 was chosen in terms of structure conservation of the X-ray structure after 50 ns of simulation. S Fig. 1A shows the comparison of the better six (out of 19) plausible protomers of the WT structural model. The best state clearly shows the lowest RMSD, which remains below 0.93 Å after 40 ns of the simulation used. In addition, the co-crystallized inhibitor perfectly conserved its structure, main contacts and conformation during the whole simulation, with RMSD below 0.38Å (S Fig. 1B.).

### 3.3 Eqolisins have four strictly conserved Glycines

Three highly conserved glycine residues, G8, G41 and G44, found in the inner sheet together with strictly conserved G55, are, given glycine’s high degree of liberty, envisaged to play a role in enzyme dynamics. EFAo000001 has G41E and G44S substitutions with surrounding subsequences substantially different from the otherwise highly conserved part of the MSA (Fig. 2). Hence, this sequence unlikely encodes for a functional eqolisin and G41 as well as G44 might be considered as strictly conserved.

In order to analyze the roles of the four glycines we performed molecular dynamics with the GAx4 mutant as compared to WT. Fig. 3A clearly shows that the GAx4 substitution yielded a much higher backbone RMSD, similar results were obtained for the overall RMSD on the inhibitor. A view of the structure averaged over the last 5 ps of the simulation (Fig. 3B1) shows major changes in the 70’s and 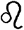-loops as well as a third loop that is stabilized by a disulfide bridge, hence referred to as the C-loop. The 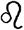-loop causes a clear loss of hydrophobic contacts with the transition state inhibitor. In addition, the GAx4 mutant shows both a repositioning of the catalytic dyad (E136/Q53) (Fig. 3B2) and a depleted H-bond donor/acceptor capacity, as compared to the WT (Fig. 3C), the net loss being between 3 and 4 H-bond contacts on the average.

**Fig. 3.**
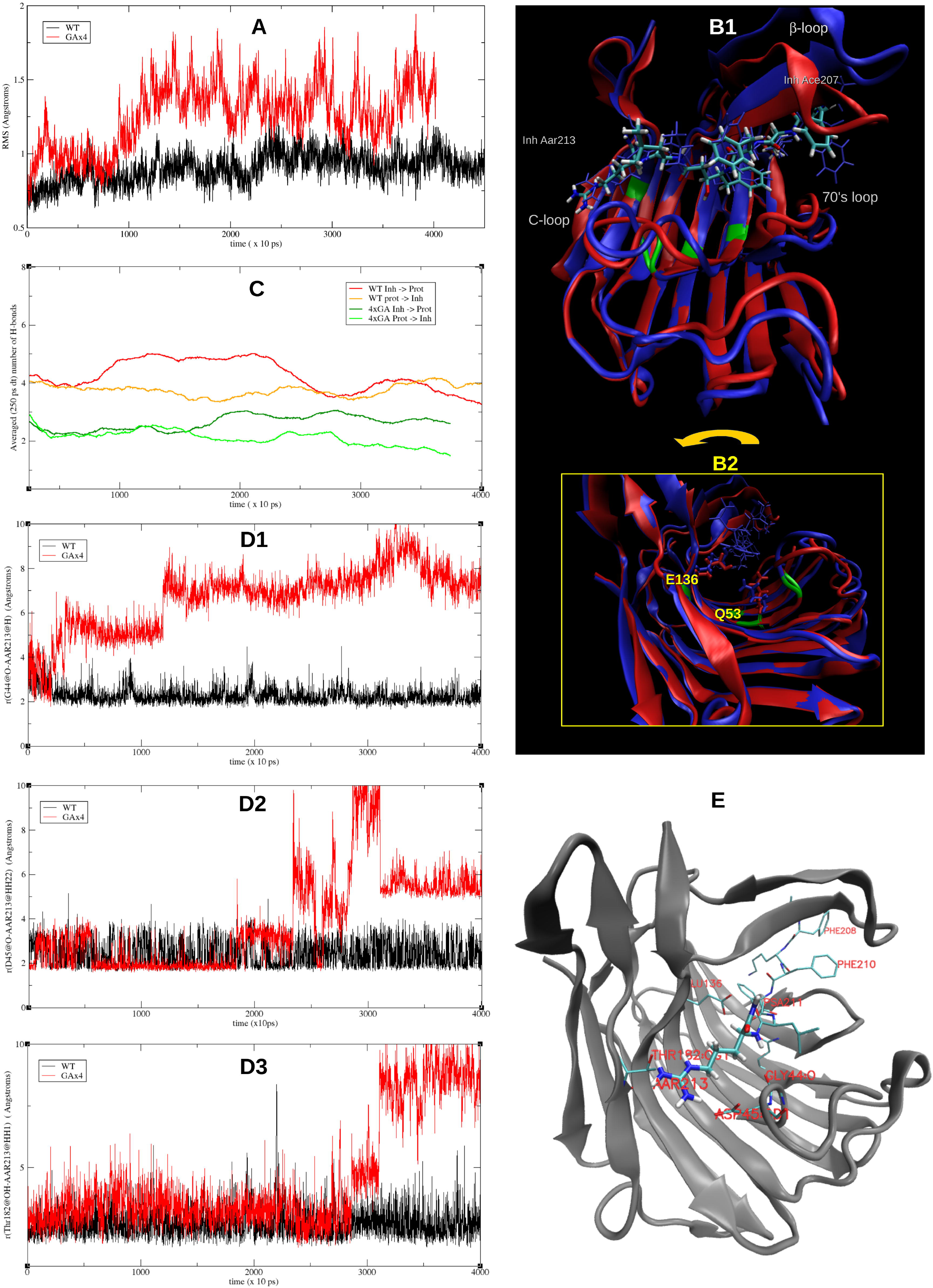
Molecular Dynamics Analyses of G8A-G41A-G44A-G55A mutant. (**A**) Comparison of the backbone RMSD of the WT and the GAx4 mutant.(**B1**) Comparison of the initial (red) and final (37.000 to 37.005 ns, in blue) structures of the GAx4 mutant. The inhibitor (Inh) represented in licorice. the mutated residues G8A-G41A-G44A-G55A in green (on the cartoon). (**B2**) Rotated view with the catalytic dyad indicated. (**C**) Number of H-bonds (smoothed as running average over 250 ps) as donor and as acceptor of the WT and the GAx4 mutant along the dynamics. (**D1**) Depletion of the contacts to the inhibitor in the GAx4 mutant (red) with respect to the WT (black) directly due to the G44A substitution, the backbone O with the backbone H of the inhibitor residue AAR213 is weakened. (**D2**) G44A substitution causes that the salt bridge involving the contiguous D45 with AAR213 is weakened. (D3) G8A substitution causes a weakened H-bond involving Q182 alcohol O with AAR213.

The above considerations preliminarily reveal a serious change in both structure and functionality. In order to confirm these observations, the actual standard free energy of binding of the eqolisin/inhibitor complexes were calculated for the WT and the GAx4 mutant using two different models (GB and PB) on the last 35 ns of the equilibrated trajectories (Table 1). The substitution of these essential glycines causes a depletion of about 33 and 34 kcal/mol (GB and PB models, respectively). An analysis of the essential modes of motion of the complex and the individual contacts was required to rationalize such a high change in the affinity for the substrate for the mutant as well as for establishing the role of each of the four G to A mutation. The overall crosscorrelation function of G41, G55 in their 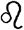-sheets and G44 (and the subsequent C-loop) shows they are correlated with the inhibitor (res # 207-213), the 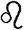-loop (res. #141-148), the 70’s loop as well as with Q53 and W67 (Data not shown), correlations that are diminished or lost in the GAx4 mutant.

**Table 1.**
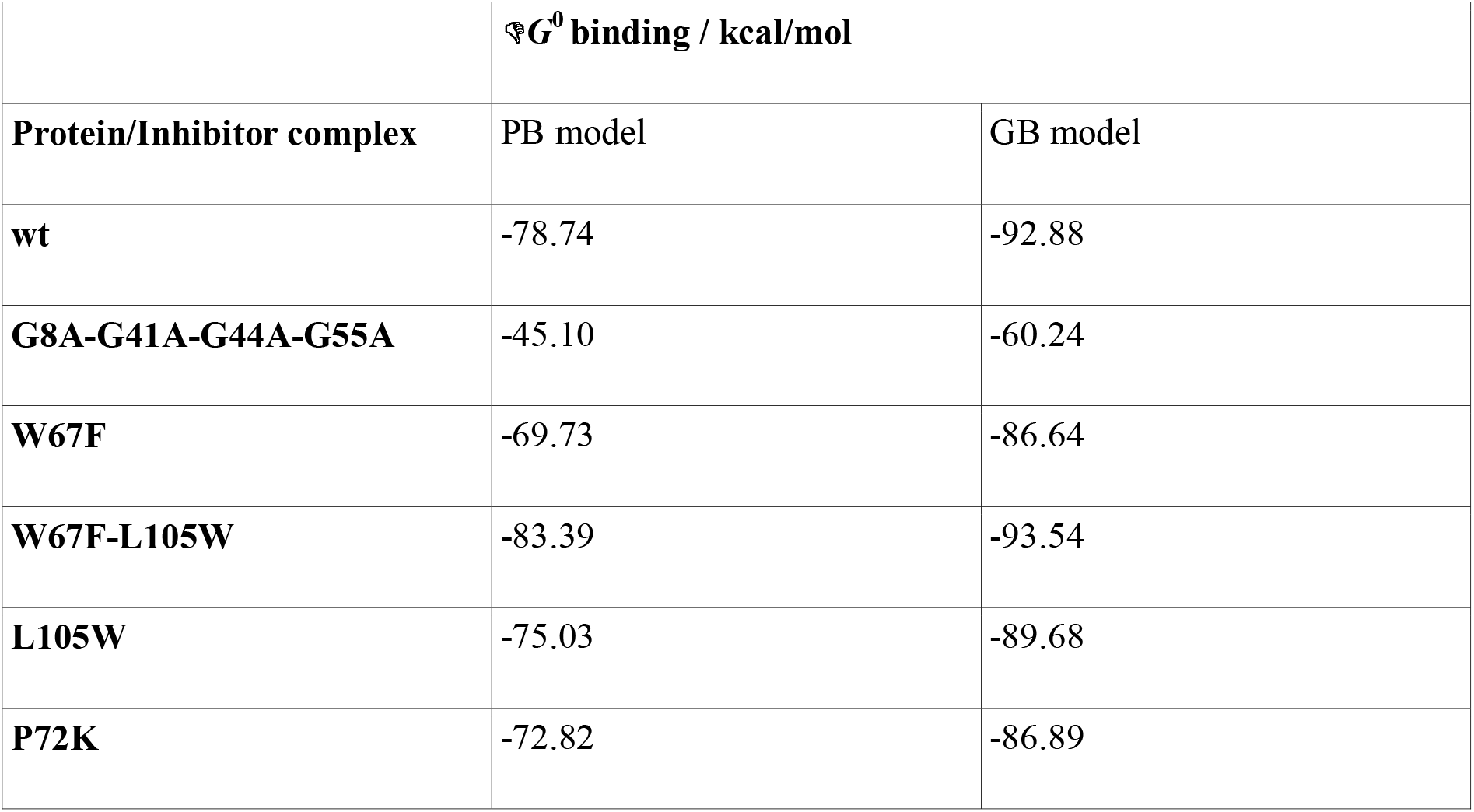
Calculated free energies of binding.

The correlations of their motions, some of which are rather long-ranged, can be rationalized by means of the contribution given by the first three lowest essential normal modes shown on S Fig. 2. Two of the three first modes are remarkably different for GAx4 with respect to WT.

Two additional evident observations about the role of these glycines are revealed in the dynamics of the WT. G44 itself is forming an H-bond to the AAR213@H residue of the inhibitor through its backbone O. The G55 is tightly H-bonded through their backbones with W67 and G44; the latter is at the beginning of the turn lead by D45, which is comprised in a tight salt bridge with the positively charged guanydonium group of the AAR213 residue of the inhibitor, also persistent during the whole simulation in the WT. On the other hand, G8 (which has also a high crosscorrelation value with AAR213 and participates into the lowest essential mode) is H-bonded with the backbone of Q182, which is also H-bonded to AAR213 through its alcohol oxygen during the whole simulation of the WT. The change in the flexibility of the backbone of G4 and G44 by the alanines in the mutant clearly affects both directly and indirectly these three strong contacts: D45@Od – AAR213@Hh22 (because of A44), A44@O–AAR213@H (because of A44 itself) and Q182@Og – AAR213@Hh12 (because of A8). Fig. 3D and E illustrate the persistence of these contacts along the whole simulation for the WT and their depletion in the GAx4 mutant.

As a result of these evaluations, we consider G8, G41 and G44 as strictly conserved, which we included in the sequence scrutiny in order to obtain a specific dataset that lacks noise in the identification of specificity determining positions (SDPs).

### 3.4 Co-evolution of W67F and L105W Mutations resulted in higher Binding Energy to Inhibitor

Another highly conserved residue is W67. Fig. 4A shows that the bulky and rigid side-chain of W67 is involved in positioning Q53, assisted by the strictly conserved E69. Furthermore, its benzene ring forms a π–π stacking with the PSA211 from the transition state inhibitor that was crystallized with SCP-B (See Fig. 4B). This indicates a role in substrate binding. A total of 15 sequences show substitution W67F. Albeit smaller, a phenylalanine can still be envisaged to fulfill both hypothesized functions. Interestingly, all sequences, although from a single clade, also show W105 which is an otherwise highly conserved small hydrophobic residue (L, I, V or M) in the outer pleated β-sheet directly below position 67. The more bulky W105 can be envisaged to change the conformation of the inner pleated β-sheet such that the aromatic ring of the F67 occurs at the same position as the typical W67, thereby sustaining for the π–π stacked stabilization of the substrate. In order to test this hypothesis and isolate the main structural factors which could rationalize it, the W67F, L105W and the double mutant (W67F/L105W) were simulated by means of molecular dynamics. The π–π stack of mutant W67F shows a similar RMSD as the WT, L105W shows a slightly higher RMSD whereas double mutant W67F/L105W showed a slightly reduced RMSD. (Fig. 4C and D). Similar results were obtained for total inhibitor RMSD (S Fig 3). Additional noticeable differences arose, especially in the case of L105W, which was unable to keep residue 209 of the inhibitor tightly bound during the whole simulation. Subtle differences, with impact in the overall energy balance, also appeared for W67F. L105W and W67F show about 3 and 6 kcal/mol, respectively (similar for PB and GB models See Table 1), less affinity for the substrate analog than the WT. On the other hand, the double mutant yielded a free energy of binding more favored than the WT (4 and 1.3 kcal/mol more negative than the WT for PB and GB, respectively). This suggest the bulkier W105 pushes F67 up to the aromatic moiety of the inhibitor, thus leading to a 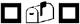 interaction even better than if it were W67/L105.

**Fig. 4.**
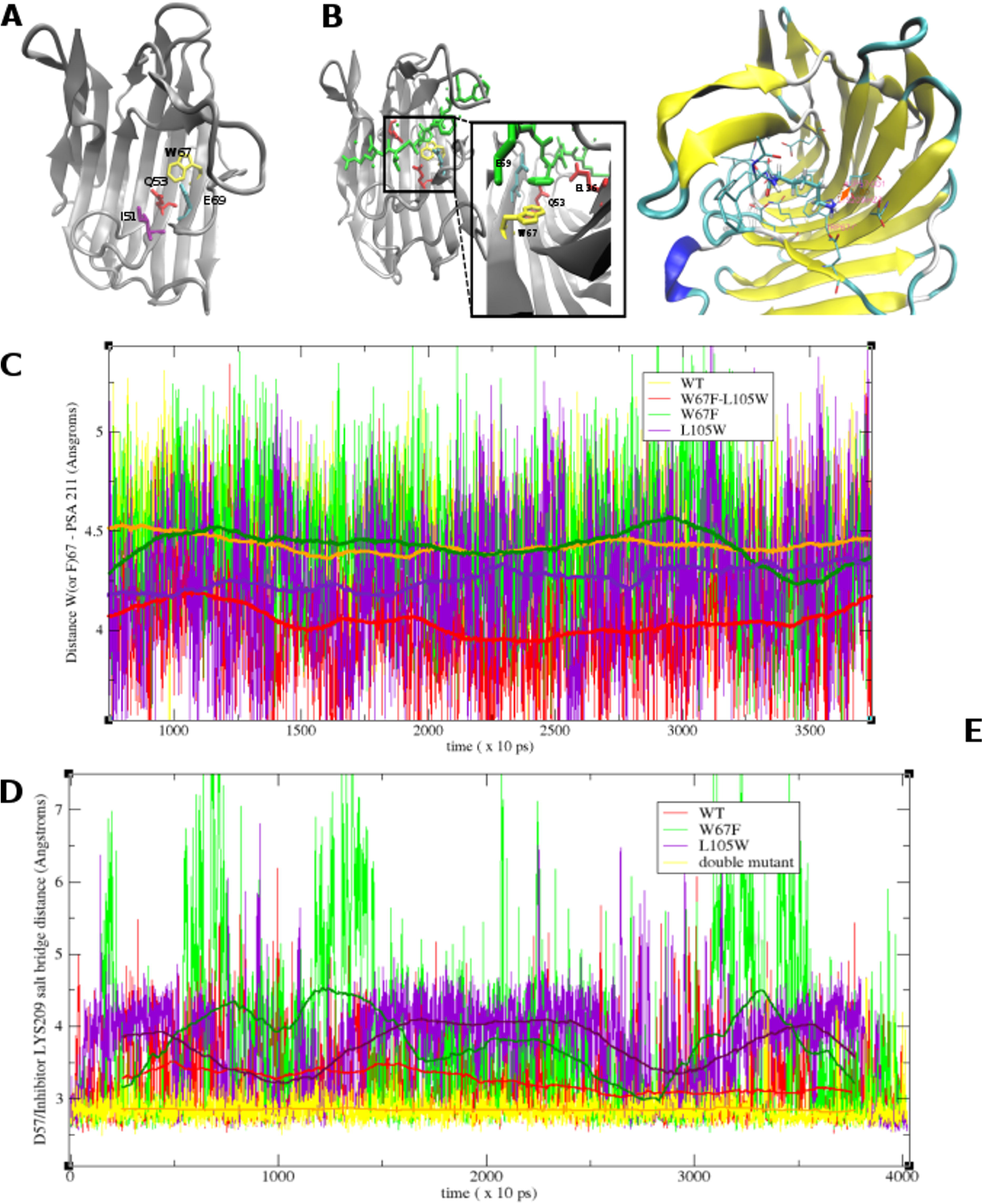
Structural and Molecular Dynamics of W67F and W67F-L105W mutants. (A) Cartoon of 2IFW showing I51 (purple) at the start of a β-sheet as well as W67 (yellow) and E69 (cyan) relative to catalytic Q53. (B) Cartoon of 2IFW detailing the π-π stack between W67 (yellow) and P2-Phe from the transition state inhibitor. (C) Cartoon of 2IFW showing 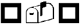 stacking (green licorice) and salt bridge (orange licorice) between eqolisin and transition state inhibitor. (D) Distance and its running average (over 500 ps, smoother solid line) for the 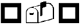 stacking of residue 67 with the inhibitor PSA211 phenyl ring in the WT, the W67F and L105W mutants as well as the W67F-L105W double mutant as determined by molecular dynamics simulation (E) Distance between D57@Og and LYS209@Hz in the WT, the W67F and L105W mutants as well as the W67F-L105W double mutant as determined by molecular dynamics simulation.

Besides its direct interaction with PSA211, W67 seems to have other important mutual interactions, which were demonstrated in the WT dynamics. The squared cross-correlation function of W67 had an important peak for PSA211, but it also was correlated with G55, D57, D65, D77, L105 and D136 (all of them >0.55, S Fig. 4). In the double mutant, a bulkier W105 instead of the L not only pushes F67 up to the inhibitor, it also repositions D57, in the WT comprised in a salt bridge interaction with the LYS209 of the inhibitor. A stable H-bond network between LYS209, D65 and D77 also depends on the right position of D57 for a tight salt bridge. Fig. 4C and 4E show the persistence and closeness of this contact during the simulation. Indeed, the combined W67F and L105W substitutions were found to be the optimal combination for properly holding D57 tightly bridged to the inhibitor. This would contribute to explain the favored binding energy for the double mutant. Besides its small improvement in the binding energy, it is worth to mention that the analysis of all the contacts (H-bonds, electrostatic and hydrophobic) from the double mutant closely resembles the analysis for the WT. Also the first essential modes are very similar to the WT as shown on the S Fig. 2. Thus whilst, either W67F or L105W substitution have little effect or deplete the binding modes and energy, the double mutant should equal or even improve the WT ability for binding this substrate/ transition state analogue. As a result of these evaluations, we consider both the W67F and the L105W as functional substitutions that seem to have co-evolved.

### 3.5 The P to K Substitution at Position 72 depletes Affinity to Inhibitor

P72 is part of the 70’s loop, described by Pillai and collaborators [14] and six sequences in the initial set of sequences have a substitution in that position. First the aforementioned NFH EFBso00001 has a serine. Then, EFPde00001 (Fig. 2), EFMac00003, EFMan00003, and EFCm000003 have a lysine whereas EFCm000002 has a glutamine. The P72K substitution was modeled and a structural alignment (data not shown) shows a structural change in the β-loop. These sequences occur at relative large distances in a preliminary tree, which is indicative for NFHs. Hence, all this suggest a proline at position 72 is required for eqolisin function.

In order to test the hypothesis that P72K is fatal, we performed molecular dynamics simulations with the P72K mutant. The structure is, in general, well conserved during the simulation (see S Fig. 5A, showing the backbone RMSD compared to the WT). The substitution has an overall effect less evident than in the case of W67F and fairly smaller than GAx4. Most contacts and the overall mode of binding to the inhibitor is similar to either the WT and the W67F-L105W. However the mobility of the 70’s loop is affected and additional interactions involving the ammonium group of K62 and alternatively N144@O N144@Od appeared, thus altering the essential mode involving the 70’s- and β-loops (see S Fig. 5B); all these interactions were absent in the WT, having the hydrophobic and compact proline residue. The substitution P72K thus depleted the affinity of the mutant for the inhibitor by about 6 kcal/mol in both models for computing the 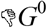 of binding. As a result of these evaluations, we consider the P72K mutation as fatal for functionality and sequences were removed accordingly. Since the P72S mutant also contains Q51 (see below) at a strict small hydrophobic site this is also pseudogene. Then, the remaining P72Q is also likely fatal. All the corresponding sequences were removed.

### 3.6 Additional low Frequency Substitutions.

A number of sequences have substitutions at position D43, which forms a minor turn in 2IFW (not shown) and has been shown to be important for function [14]. Three sequences, one bacterial and two eukaryotic, have S, which, when phosphorylated, is physicochemically similar to aspartic acid, hence these are envisaged non-fatal substitutions. Ten sequences have G43, a relatively large number that, combined with its appearance in fungi, bacteria and archaea suggests this is another acceptable substitution. Furthermore G is, given its high degree of liberty, often found at turns.

Position 51 has a highly conserved small hydrophobic residue, L, I or V, where EFTa000006 has P instead. The isoleucine in 2IFW occurs at the start of an inner sheet element and has its side-chain pointing inward the halfpipe (Fig. 3A). It seems unlikely that substitution by the rigid proline would not affect catalytic efficiency, hence we consider EFTa000006 (Fig. 2) as an NFH. EFBso00001 has Q51 (not shown) as well as the aforementioned P72Q mutation, hence this sequence also likely encodes an NFH. As such it seems a, likely small, hydrophobic residue is required at position 51. Sequences were removed accordingly.

Three bacterial sequences have a phenylalanine at position 89 that is otherwise characterized by VILM. Since phenylalanine is hydrophobic, this appears as an allowed substitution. Position 105 is another conserved hydrophobic site, typically constituted by VILM but shows, besides the above discussed Ws, two instances of phenylalanine, one of tyrosine and one of glutamate. The glutamate substitution in BCa0000005 would result in a buried but charged side-chain, which could be explained by a co-evolved basic residue, which could not not identified. Hence, this is likely a fatal substitution. The tyrosine substitution in APf0000001 might well be a problem of erroneous sequencing or gene modeling since it is surrounded by a stretch of about 20 to 40 amino acids that poorly align and the sequence was removed. The two instances of phenylalanine substitutions in bacterial (Bgt0000001) and eukaryotic EFCm000005 introduce another hydrophobic but larger residue and given the partial solvent exposure, it can be envisaged this substitution is not fatal. Finally, the highly conserved site 133 has VILM and two, likely partial sequences (EFFo000002 and EFNh000001) showing a gap in the MSA were also removed.

### 3.7 Phylogenetic Clustering suggests Fungal Eqolisins have resulted from an ancient LGT Event.

Following the sequence scrutiny an additional 23 scrutinized eqolisin sequences from leotiomycetes were added, a new MSA of a total of 315 sequences was made (S datafile 2), subjected to trimming of poorly aligned subsequences and used for phylogeny. First a maximum-likelihood tree was constructed, which was then used to initiate a Bayesian analysis. Fig. 5A shows an annotated radial phylogram, a radial cladogram is shown in Fig. 5B. Cluster assignation was made based on monophyly and, more importantly, taxonomical considerations. Apart from two orphan sequences, prokaryotic (archaeal and bacterial) and fungal sequences cluster separately which suggests a common ancestor, separating prokaryotic clades I and II from the fungal sequences. However, a taxonomical distribution analysis (Fig. 5C) points towards an alternative common ancestor, also indicated in Fig. 5A. The majority of the sequences are found in the subphylum pezizomycotina, which is further classified in the classes of dothideomycetes, eurotiomycetes, leotiomycetes, sordariomycetes and pezizomycetes, the last class not containing eqolisins. Clade III is a monophyletic clade with sequences of all four classes and since it is closest to the initially suggested common ancestor, this indicates the common ancestor might correspond better with the node between the clade containing subclades I, II and III and the rest of the tree. This is substantiated by the sequence logo analysis (Fig. 6A) presented in the description of the structure function analysis. All other assigned clades were selected based on monohyly and size (a requirement of at least 10 members was set arbitrarily).

**Fig. 5:**
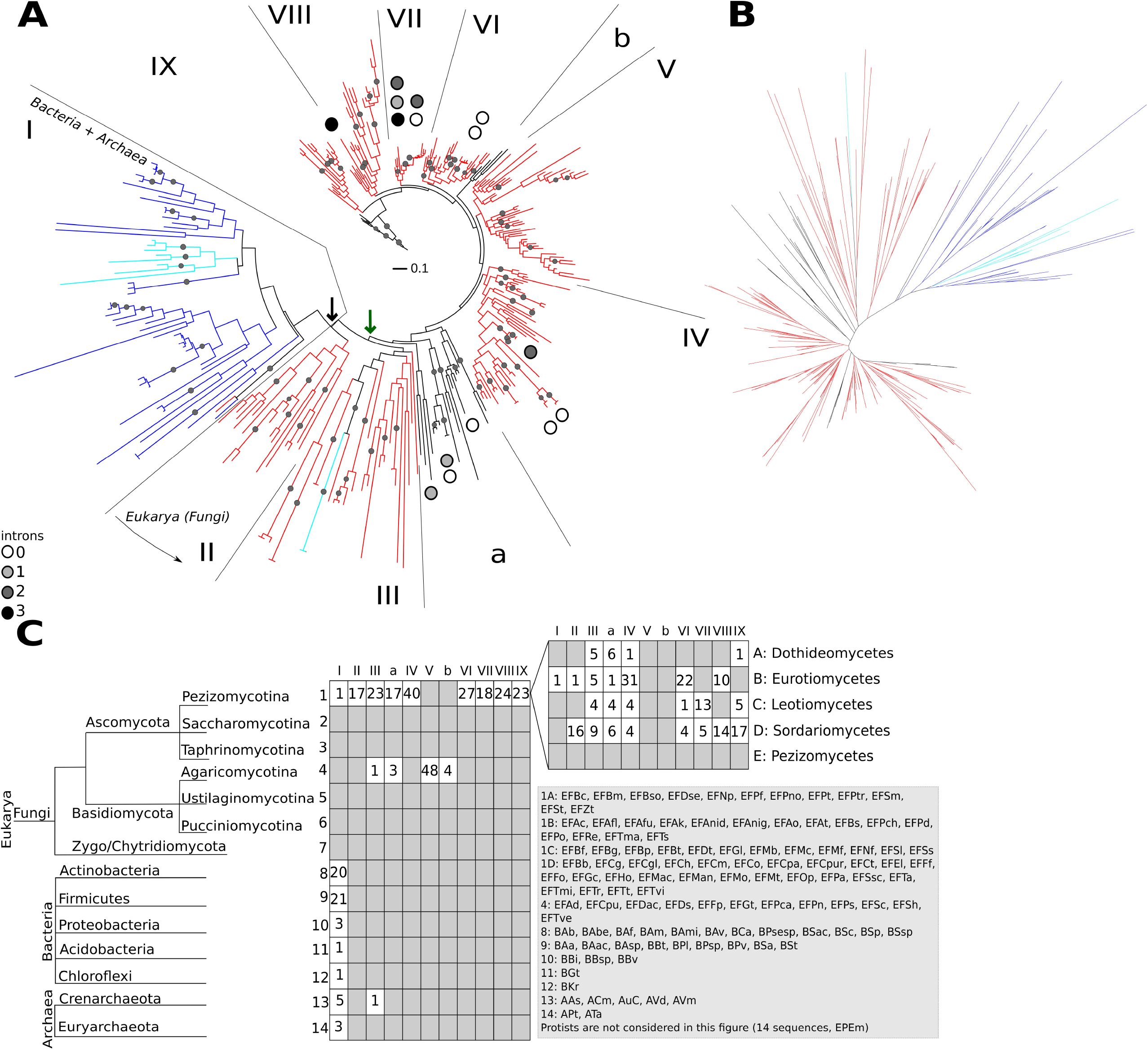
Phylogenetic Analysis of Eqolisins. (A) Radial phylogram. Fungal sequences are in red or black (small clusters indicated with a and b), Bacterial sequence in blue and Archaeal in cyan. The black and green arrows indicate two possible common ancestors discussed in the text. Roman numerals refer to selected clades discussed in the main text. Dots indicate Bayesian support > 0.8, scale bar represents a distance of 0.1 accepted amino acid mutations per site. (B) Radial cladogram. (C) Taxonomic distribution of eqolisins on a tree representative for generally accepted consensus species phylogenies. Roman numerals and a and b refer to the selected clades (Fig. 5A). Numbers indicate the number of sequences per clade and subphylum/class. Abbreviations in S datafile 3.

**Fig. 6.**
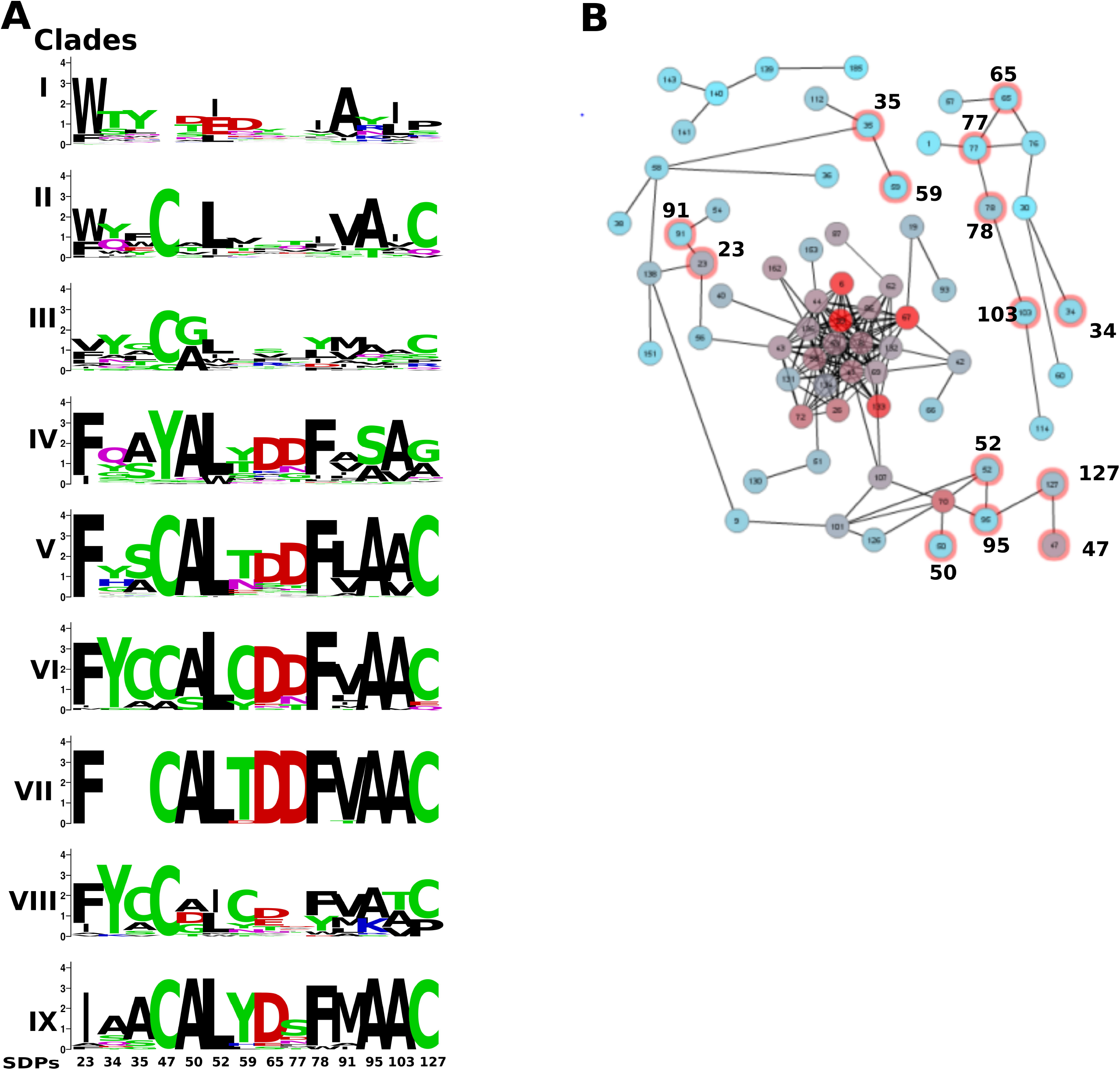
Specificity Determining Positions of Eqolisins. (A) Sequence logos of SDPs per selected clade (Fig. 5). (B) Mutual Information Network. Connected nodes have MI of at least 6.5. Note that a first SDN (SDPs 47, 52, 95, 127 and 50) is connected to the major network, whereas the other SDN ((65, 77, 78 and 103) is. Colour of the nodes corresponds to Kullback Leibler conservation.

Lateral gene transfer (LGT) seems to have had an important role in the eqolisin phylogeny. The two orphan sequences are clear examples of LGT. Furthermore, the lack of eqolisin encoding sequences in eukaryotes other than fungi points towards an ancestral LGT event. The taxonomical distribution of sequences suggests this ancestral LGT has taken place between a bacterium and the ancestor of the Dikarya. The absence of introns in many, albeit not all fungal sequences corresponds with the proposed ancestral LGT

### 3.8 Functional Diversification of Substrate Binding Site

Since a number of fungi have various paralogs in well separated clades, fungal eqolisins might have been subject to functional diversification. We performed analyses in order to identify first Cluster Determining Positions (CDPs) and then Specificity Determining Positions (SDPs). CDPs are positions in the protein (or columns in the corresponding MSA) that significantly contribute to clustering. These are the result of either genetic drift (i.e. neutral substitutions) or selection, which would mean they are somehow related to functional or structural diversification. Functionally, but also structurally, important residues are likely to show moderate to high levels of interaction with other residues, which can be determined with the measure of mutual information (MI). Hence, CDPs that show high MI with other residues are likely important in either maintaining the structure or functional diversification. Since MI reflects co-variation, groups of CDPs and other positions that are directly connected via significant MI values are expected to affect the same functional aspect.

CDPs were identified using SDPfox [40], using the clustering indicated in Fig. 5. Out of 28 CDPs that were identified, 20 showed significant cumulative MI levels, as determined by Mistic [41], of which a total of 14 could be confirmed by H2Rs [42], all considered putative SDPs (pSDPs). A resume of the SDP identification is shown in S Datafile 4 that also contains a description of the binding cleft, based on the work of Pilai and co-workers on SCP-B [14]. Sequence logos of the SDPs, according to the clustering shown in the phylogeny of Fig. 5., are shown in Fig. 6A alongside the MI network, defined as the subnetwork of nodes that directly connect to at least one other node with significant MI, obtained by Mistic (Fig. 6B). The MI network consists of three fully connected subnetworks. The large connected subnetwork contains an intricate central module of nodes, mostly corresponding to highly conserved sites, with some branches with lower levels of connections. This central module conceptually corresponds with the core eqolisin function and lacks SDPs. Two instances of paired SDPs, (35 and 59; and 23 and 91 respectively) are found in one of the branches with low connection levels and appear to be the result of co-evolution driven by of structural compensation. The sequence logo shows SDP35 and 59 have a preferred C in both clade VI and VIII that, combined with their proximal 3 dimensional location, points toward a possible disulfide bridge. SDPs 23 and 91 are likely involved in an important hydrophobic interaction given the substitution pattern shown in Fig. 6A and their 3 dimensional proximity in the protein (not shown). These pSDPs are as such not considered real SDPs since we cannot foresee any functional diversification. A first specificity determining network (SDN) with four directly connected SDPs (47, 52, 95 and 127) as well as SDP50 that connects via single non-SDP to SDP52 and SDP95, is found in one of the branches with lower levels of connections. A second SDN with four directly connected SDPs 65, 77, 78 and 103 locates into one of the small (n=11) subnetworks that also includes pSDP34. As such, the isolation of pSDP34 suggests this at least an important SDP. The second small (n=5) subnetwork contains no SDPs.

The SDN in the small subnetwork is dominated by the clearly preferred D65, D77 and F78 in groups IV to IX (Fig. 6A). Fig. 7 shows how D65 and D77, part of subsite S3, interact with the substrate. Both acid side chains interact directly with the basic sidechain of the Lys in the substrate analogue. F78 does not form part of the binding cleft, but interacts directly with its neighbor D77. A103 connects to F78, having an MI value of 6.7 and Fig. 7B shows they interact physically. Hence, SDPs 65 and 77 are predicted to affect substrate specificity, particularly concerning P3. Positions 78, 103 likely give some sort of structural support. The information contents at positions 65 and 77 in the various clades is likely indicative for substrate specificity.

**Fig. 7.**
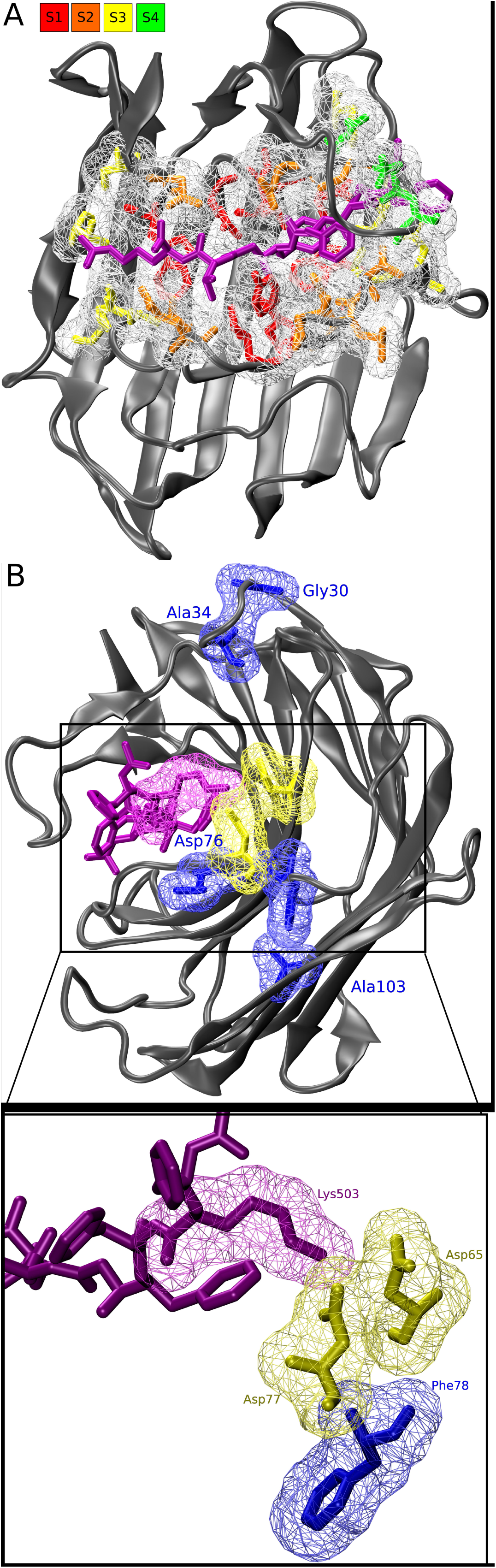
The D65-D77 Mutual Information network is involved in substrate specificity. (A) Cartoon of 2IFW with highlighted binding cleft. Transition state inhibitor is indicated in purple licorice. Sites are indicated in colored licorice (see legend) with surface indication. (B) Interaction of D65 and D77 with substrate analog. The inset shows that the acid groups of both aspartates (S3’ yellow surf) interact directly with the basic group of the Lysine form the the analog (purple). Other SDPs of the same SDN are indicated in blue.

SDPs of the same SDN are indicated in blue. The SDP group in the large subnetwork contains SDPs dominated by the Cysteins at 47 and 127 that form a disulfide bridge in SCP-B and which are present in most sequences, except for those in clades I and IV (Fig. 6A). In SCP-B the disulfide bridge stabilizes a hitherto un-described loop from position 44 to 51 which we refer to as the C-loop. Fig. 8A shows the C-loop, the disulfide bridge as well as the 70s loop since the central position in group 2 is taken by position 70, a Tyr in SCP-B. This suggests the C-loop interacts with the 70s-loop, which has been shown to be important in binding dynamics. Fig. 8B demonstrates that in SCP-B both SDPs A50 and L52 interact physically with Y70, likely influencing the mutual positioning of the 70s- and C-loops. Clade IV sequences have Y47, which appears to be compensated by the substitution C127A. SDP127 is connected with SDP95, which is encountered in the outer sheet just below the inner sheet and the C-loop. This suggests a secondary, structural compensation but the substitution pattern (A95S) does not give any clue on how this might be established. The C-loop contains four residues that are part of the binding cleft (Supplemental Table 1) of which, interestingly 51 is another CDP that does not form part of the network. All together this suggests that the C-loop is involved in the dynamics that lead to substrate binding.

**Fig. 8.**
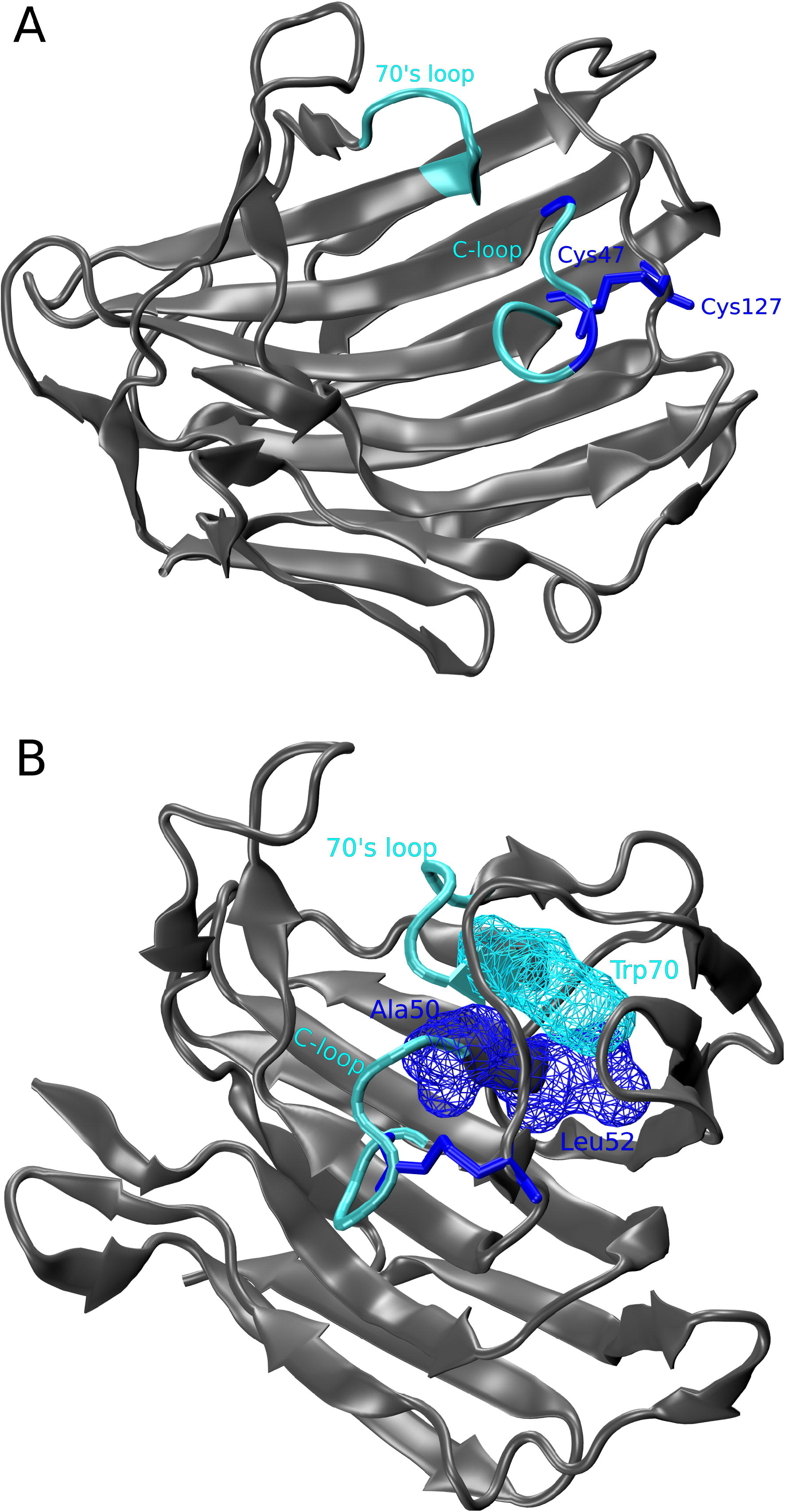
The C-loop interact with the previously described 70s-loop. (A) Cartoon of 2IFW with highlighted the previously described 70s-loop and the novel C-loop (Cyan). C47 and C127 (blue licorice) form a disulphide bridge that holds the loop inplace. The C-loop also includes SDP50 (blue). (B) Cartoon of 2IFW with highlighted the 70s- and C-loop. Y70 interacts physically with A50 and L52, both identified as SDPs

## 4 Conclusions

Only few eqolisins exist but interestingly fungi that secrete acid as part of their lifestyle can have up to nine paralogs, which implies functional redundancy and diversification. This was studied using a stringently mined sequence set. A number of sequences we removed might be functional but we prefer to prevent possible contamination with sequences of NFHs. An interesting case of recent molecular evolution was identified, resulting in a most likely more active enzyme. Furthermore we detected two groups of SDPs, one that very likely is involved in substrate specificity given two of the SDPs form part of the binding cleft. The other group seems to affect loop dynamics. Some of the additionally identified CDPs might also be SDPs but the lack of MI signal cannot corroborate them.

## Legends Supplemental Figures

**S Fig. 1. Molecular Dynamics Simulation determining best protomer state.** (A) Backbone RMSD of the best protomer for the WT compared to five of the other best. (B) Backbone RMSD of substrate analog inhibitor for the best protomer for the WT compared to five of the other best.

S Fig. 2. A1-A3, Ca vectors of the first essential normal mode from PCA analysis for the GAx4, WT and W67F-L105W species. A4, the squared displacement of each residue in the first mode. B-C squared displacement of each residue on the next two modes.

S Fig. 3. RMSD of the inhibitor residues for the WT, W67F, L105W and the double mutant.

S Fig 4. Squared cross correlation function of W67 against all other residues (wt trajectory).

S Fig. 5. Backbone RMSD comparison for the WT and the P72K mutant.

w1: https://www.ebi.ac.uk/Tools/hmmer/search/hmmsearch 09/04/17

w2: http://prody.csb.pitt.edu/ 09/04/17

